# Diagnosis of central serous chorioretinopathy by deep learning analysis of *en face* images of choroidal vasculature

**DOI:** 10.1101/2020.12.11.421040

**Authors:** Yukihiro Aoyama, Ichiro Maruko, Taizo Kawano, Tatsuro Yokoyama, Yuki Ogawa, Ruka Maruko, Tomohiro Iida

**Author notes:** Corresponding author: Ichiro Maruko, MD, Department of Ophthalmology, Tokyo Women’s Medical University School of Medicine, Shinjuku, Tokyo, Japan, (IM). All authors contributed equally to this work. Dr. Aoyama has nothing to disclose. Dr. I. Maruko reports grants from JSPS KAKENHI (Grant Number JP20K09781), grants and personal fees from Alcon Pharma K.K., personal fees from Bayer Yakuhin, Ltd., personal fees from Santen Pharmaceutical Inc., personal fees from Alcon Japan, Ltd., personal fees from Topcon Co., Ltd., personal fees from Senju Pharmaceutical Co., Ltd., personal fees from NIDEK Co., Ltd., outside the submitted work. Dr. Kawano has nothing to disclose. Dr. Yokoyama has nothing to disclose. Dr. Ogawa has nothing to disclose. Dr. R. Maruko has nothing to disclose. Dr. Iida reports grants and personal fees from Alcon Pharma K.K. (Japan), personal fees from Bayer Yakuhin, Ltd. (Japan), grants and personal fees from Santen Pharmaceutical Co., Ltd. (japan), grants from Nidek (Japan), grants from Senju Seiyaku (Japan), research support from Canon (Japan), research support from Kowa (Japan), research support from Topcon (Japan), outside the submitted work.

## Abstract

**Purpose:** To classify central serous chorioretinopathy (CSC) by deep learning (DL) analyses of *en face* images of choroidal vasculature obtained by optical coherence tomography (OCT) and to analyze the regions of interest for DL from heatmaps.

**Methods:** One-hundred eyes were studied; 53 eyes with CSC and 47 normal eyes. Volume scans of 12×12 mm square were obtained at the same time as the OCT angiographic (OCTA) scans (Plex Elite 9000 Swept-Source OCT^®^, Zeiss). High-quality *en face* images of the choroidal vasculature of the segmentation slab of one-half of the subfoveal choroidal thickness were created for the analyses. The entire 100 *en face* images were divided into 80 for training (100 times) and 20 for validation. The Neural Network Console (NNC) developed by Sony and the Keras-Tensorflow backend developed by Google were used as the software for the classification with 16 layers of convolutional neural networks. The active region of the heatmap based on the feature quantity extracted by DL was also evaluated as the percentages with gradient-weighted class activation mapping implementation in Keras.

**Results:** In the 20 eyes used for validation including 8 eyes with CSC, the accuracy rate of the validation was 100% (20/20) for NNC and 95% (19/20) for Keras. This difference was not significant (*P*=0.33). The mean active region in the heatmap image was 12.5% in CSC eyes which was significantly lower than the 79.8% in normal eyes (*P*<0.01).

**Conclusions:** CSC can be automatically classified with high accuracy from *en face* images of the choroidal vasculature by DLs with different programs, convolutional layer structures, and small data sets. Heatmap analyses showed that DL focused on the area occupied by the choroidal vessels and their uniformity. We conclude that DL can help in the diagnosis of CSC.

## Introduction

Artificial intelligence (AI) or machine learning using deep learning (DL) techniques has achieved human-like or even beyond human performances especially in visual recognition. In Go and Shogi board games, AI is comparable to or has surpassed top-level human players, and it has led to the creation of new moves that humans could not imagine.[1–3] Despite such remarkable achievements, one of the problems of AI or DL is its lack of transparency. Because the contents of DL is a black box, it is difficult to completely explain the reasons used for the choices made even by human experts which raises concerns about accountability and responsibility.

In the field of ophthalmology, AI has been applied to the diagnosis and/or staging of diabetic retinopathy,[4–6] glaucoma,[7–9] age-related macular degeneration,[10,11] retinopathy of prematurity[12–14], and other retinochoroidal disorders.[15–17] Most of the AI methods enabled ophthalmologists to diagnose and determine the stage of the disorders with the same or slightly higher accuracy than specialists. However, where to find and produce the results include some undetermined factors.

Central serous chorioretinopathy (CSC) is a chorioretinal disorder that is associated with a serous retinal detachment in the macular region including the fovea which then leads to visual impairments.[18] The primary change is in the choroid where there is a choroidal thickening and a dilation of the large choroidal blood vessels in the middle or Haller’s layer.[19–24] We have reported that the choroidal vascular density in one-half of the choroid is higher in eyes with CSC than normal eyes using *en face* OCT images.[25] However, it was difficult to diagnose CSC based on only the choroidal vascular density.

The purpose of this study was to determine whether the choroidal vascular pattern in the OCTA *en face* images is different in CSC eyes from that of normal eyes using DL. In addition, we determined the regions of the choroidal vasculature that were used by DL to make the differentiations.

## Methods

The medical record of 100 eyes of 100 patients who had been examined by OCT and OCTA in the Department of Ophthalmology of the Tokyo Women’s Medical University between 2017 and 2018 were reviewed. The procedures used were approved the Institutional Review Board of the Tokyo Women’s Medical University School of Medicine, and they conformed to the tenets of the Declaration of Helsinki (approval number 2636-R). All of the examinations were performed after an informed consent was obtained. In our department, OCT and OCTA devices are used to study eyes with macular and retinal disorders, and observational studies of CSC and age-related macular degeneration.

Fifty-three eyes of 49 patients (39 men and 10 women) with CSC were studied. Their average age was 51.7 ± 12.7 years, and all had been diagnosed with CSC by fluorescein angiography (FA) and indocyanine green angiography (ICGA). Eyes treated with photodynamic therapy were excluded because of choroidal changes after photodynamic therapy.[26–29] Forty-seven eyes of 47 individuals (11 men, 36 women; average age 37.5 ± 5.4 years) without retinal and choroidal disorders were examined in the same way, and their data were compared to those of the CSC patients.

### Obtaining scanning OCTA and OCT *en face* images

All eyes were examined by swept source OCTA (SS-OCTA; AngioPlex Elite 9000, Zeiss, Germany) whose light source emission was between 1040 and 1060 nm. The SS-OCTA cannot obtain images of the choroidal blood flow pattern in normal eyes because of the attenuation of the observation light by the RPE. On the other hand, high quality OCT *en face* images can be obtained at the same time as the OCTA scanning. The AngioPlex Elite 9000 with 100,000 A-scans/second and a real-time eye tracking system (FastTrac™) can record 12 × 12-mm OCTA images and structured *en face* images at the same time. The standard high-resolution OCT *en face* images were used for the measurements. The segmentation boundaries and widths were manually adjusted to the choroidal areas to be analyzed by the embedded software. Only images with signal intensities of 8 or more were used to maintain the reliability of the analyses.

### Measurements of choroidal thickness

The AngioPlex Elite 9000 can record 500 horizontal 12 mm cross sectional scans with the OCTA/*en face* OCT scanning procedures. The subfoveal choroidal thickness (SCT) through the fovea was measured using the caliper tool in the embedded OCT software. The choroidal thickness was defined as the distance between the basal border of the RPE and the chorioscleral junction.

### OCT *en face* images of choroidal vasculature

The standard *en face* images were flattened at Bruch membrane to quantify the thicknesses of the different structures of the choroid. The segmentation slab selected for the analyses was set at one-half of the subfoveal choroidal thickness with a 30 μm width for the analyses as reported (Fig 1).[25] After segmentation, the image was cropped to a square and adjusted so that the center of one side of the square was at the center of the temporal edge of the optic disc. This excluded the information of the optic disc from the analyses. When peripapillary atrophy, observed as a whitish area around the optic disc, was present on the temporal side of the optic disc, the area to be analyzed was further cropped to remove the atrophic areas.

**Fig 1.**
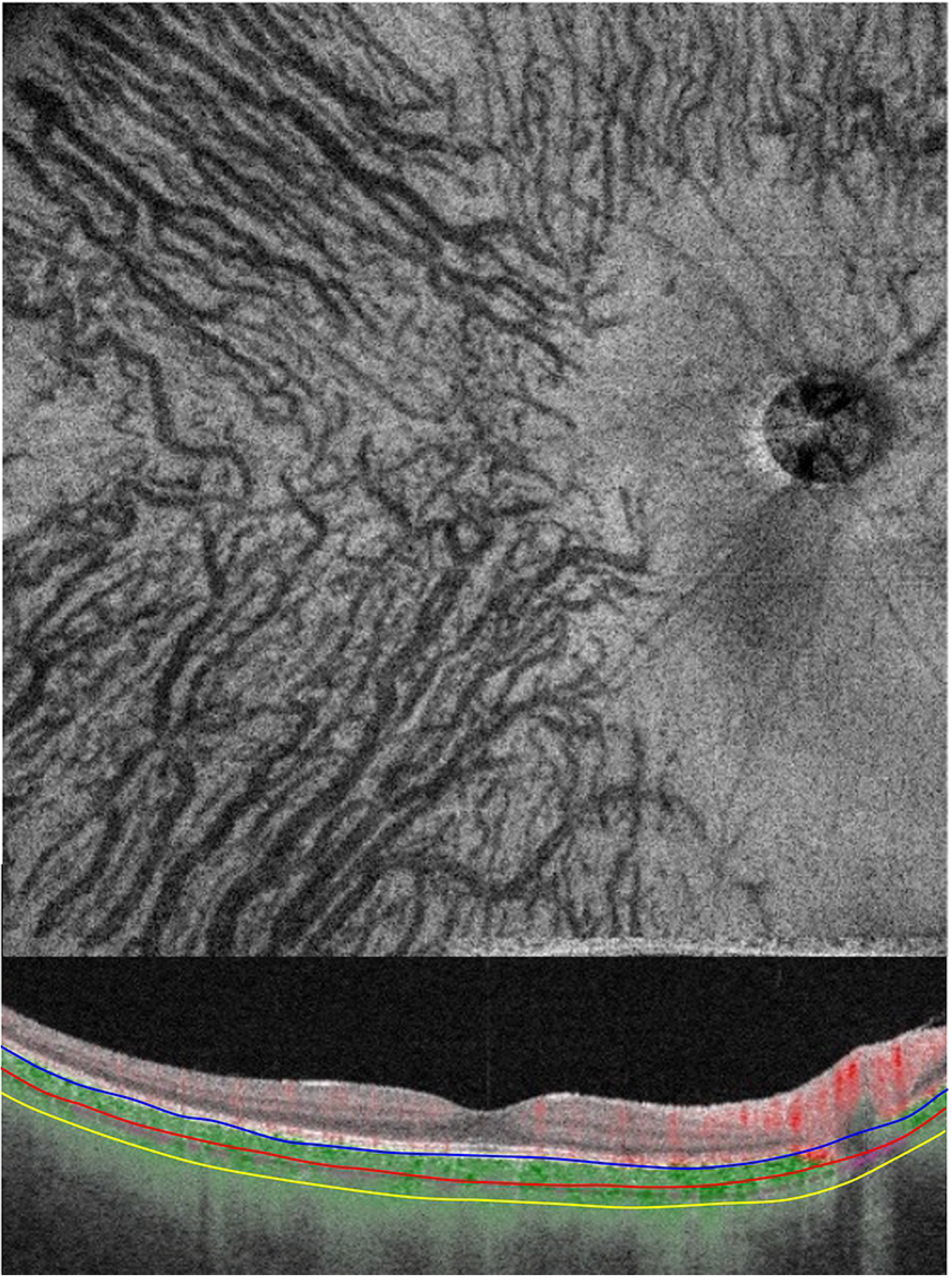
High-quality vascular *en face* image of the choroidal vascular structure at the selected segmentation slab of one-half of the subfoveal choroidal thickness.

### Deep learning

We used two Convolutional Neural Network (CNN) models: the Neural Network Console (NNC; SONY), and the Keras-Tensorflow backend (Google). The VGG16 model is comprised of five blocks with three fully connected layers. Each block includes the convolutional layers followed by a max-pooling layer. A flattening of the output from block 5 results in two fully connected layers. The NNC model resembles VGG16. The Keras model - VGG16 model is fine-tuned by transfer learning.

The original *en face* image (550 x 550 pixels) was cropped to remove the optic disc and resized into 224X224 pixel. The cropped and resized image was used for the both models. The entire 100 *en face* images were randomly divided into 80 for training (100 times) and 20 for validation. We calculated the correct answer rate during the validation process.

### Heatmap with Gradient-weighted Class Activation Mapping (Grad-CAM) implemented in Keras

Determining the characteristics of all the convoluted layers that we analyzed with Grad-CAM implemented in Keras were evaluated as heatmaps of the areas that were analyzed by DL.[30–32] The heatmap images were superimposed on the choroidal OCT *en face* vascular images to determine where the DL system was focusing its attention to in the choroid. The heatmap images were also exported to ImageJ, and the active region was calculated as a percentage of the cropped image using a threshold value of 128 out of 255 gradations in the heatmap image generated by Grad-CAM.

### Statistical analyses

All *P*-values are two-sided, and *P* values <0.05 were considered statically significant. All statistical analyses were performed with EZR free software (Saitama Medical Center, Jichi Medical University, Saitama, Japan), which is a graphical user interface for R (The R Foundation for Statistical Computing, Vienna, Austria).[33] More exactly, it is a modified version of R commander designed to add statistical functions that are frequently used in biostatistics.

## Results

Fifty-three eyes (26 right, 27 left) of 49 patients with CSC and 47 eyes (23 right, 24 left eyes) of 47 normal individuals without any retinal and choroidal disorders were studied. The mean age was 51.7 ± 12.7 years in the CSC group which was significantly older than the 37.5 ± 5.4 in the normal group (*P* <0.01). The subfoveal choroidal thickness (SCT) was 480 ± 92 μm in the CSC group which was significantly thicker than the 292 ± 64 μm in the normal group (*P* <0.01). Representative cases are shown in Fig 2 through 4.

**Fig 2.**
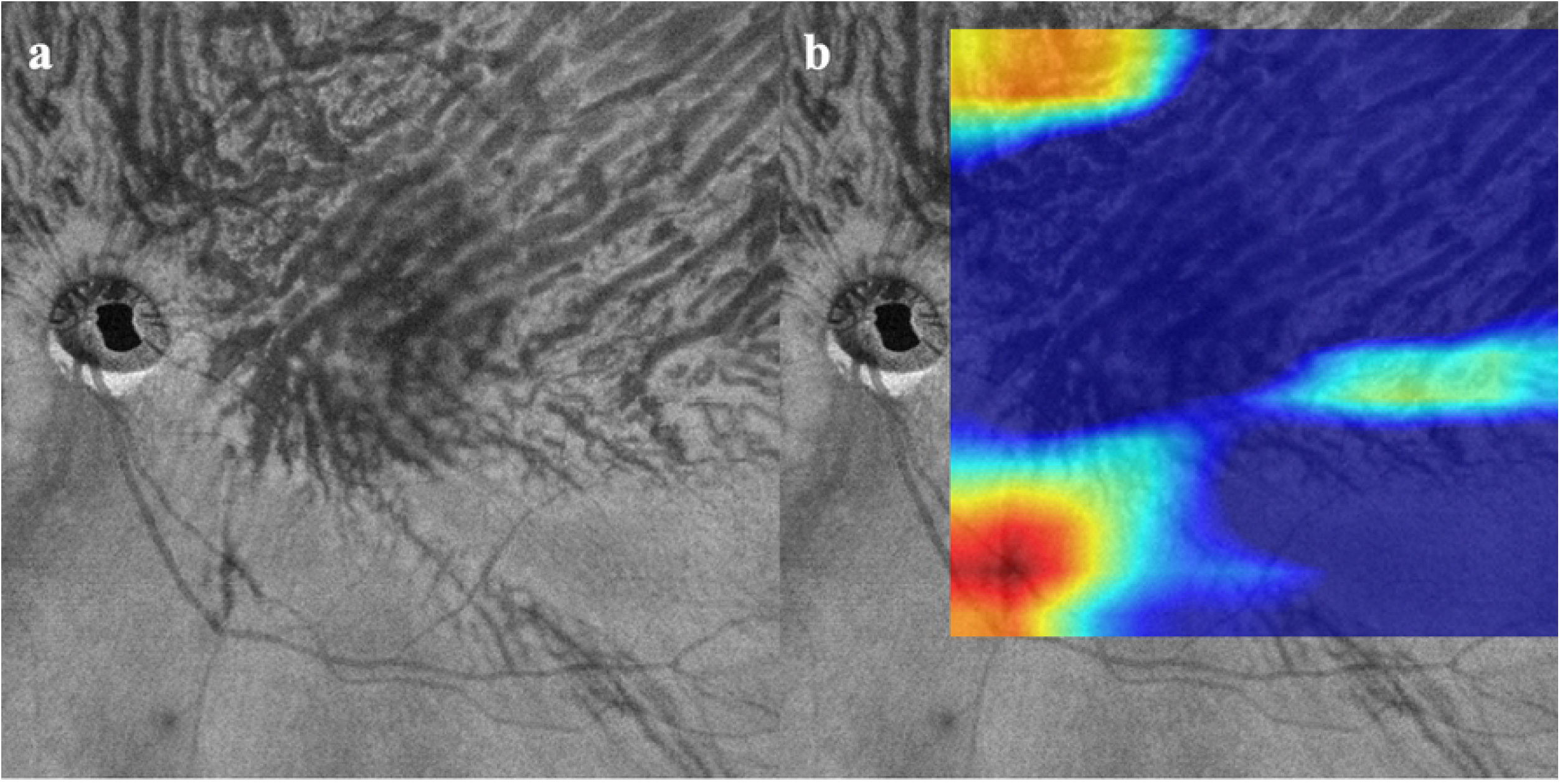
Central serous chorioretinopathy (CSC). a. Optical coherence tomography (OCT) *en face* image of choroidal vasculature. Dilatated choroidal vessels have an asymmetrical pattern flowing to the superior sector. Both the Convolutional Neural Network models of NNC and Keras determined that this image was obtained from an eye with CSC. b. Heatmep superimposed on OCT *en face* image. Deep learning did not focus on the large and dilatated choroidal vessels. The active region was 11.26%.

**Fig 3.**
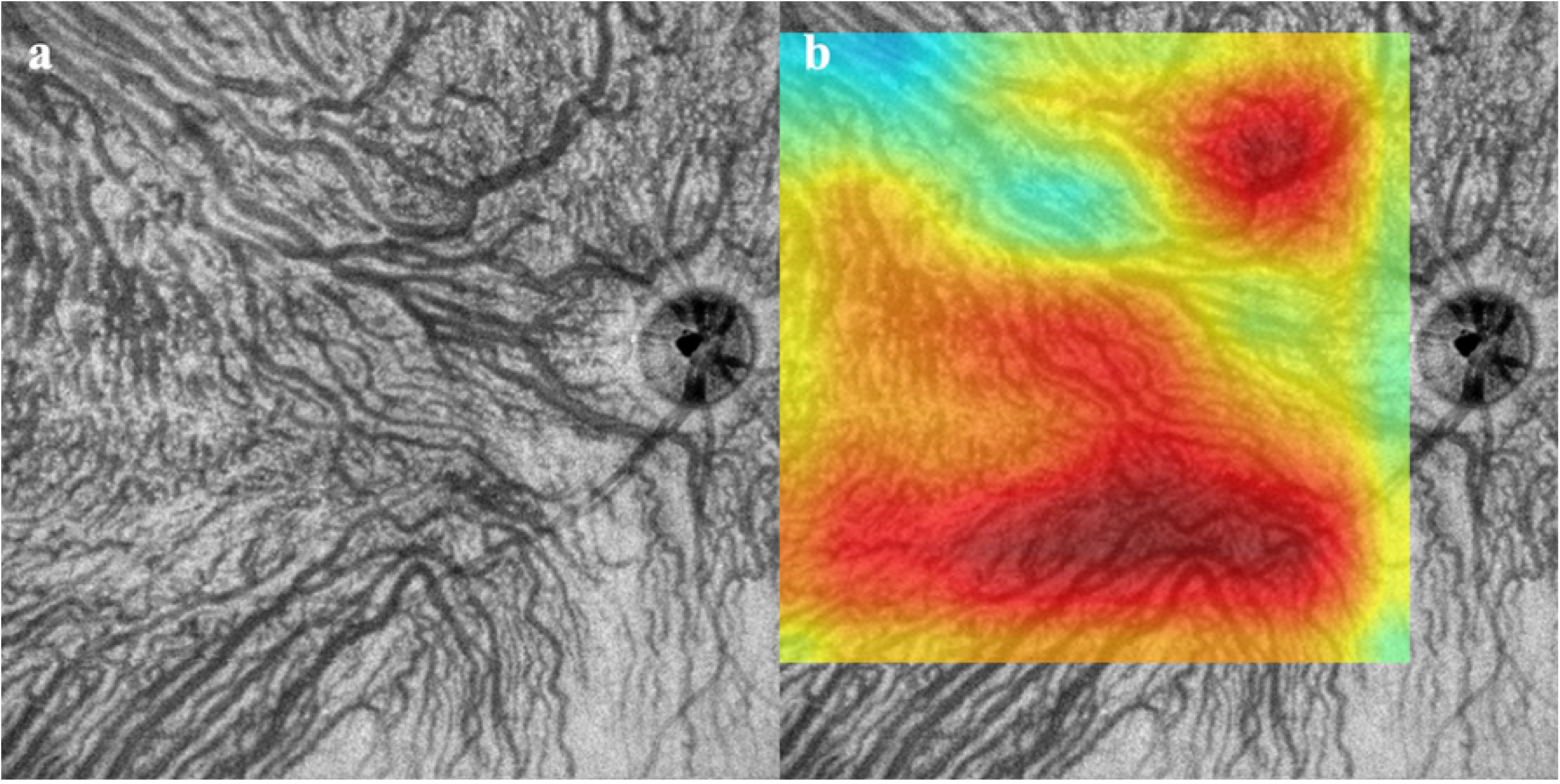
Normal. a. Choroidal vessels are distributed in a symmetrical pattern from superior and inferior sectors. Both Convolutional Neural Network models determined that this was obtained from a normal eye. b. Heatmep superimposed OCT *en face* image. Deep learning focused on the uniform choroidal vessels and the area occupied by choroidal vessels in the image. The active region was 92.22%.

**Fig 4.**
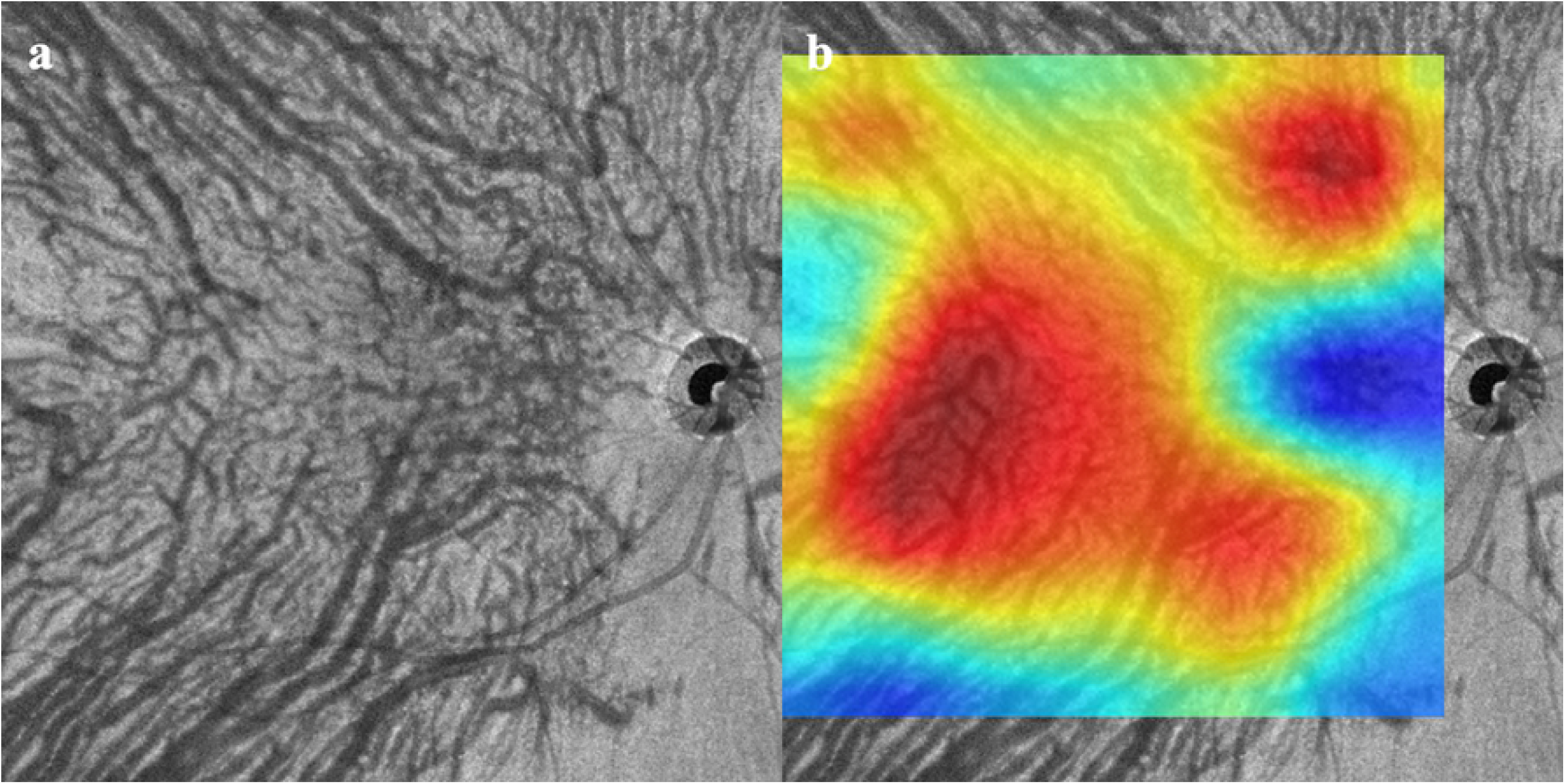
Central serous chorioretinopathy (CSC). Some choroidal vessels are dilated, and the vascular density is high. However, the choroidal vessels are distributed in vertically symmetrical pattern. The NNC model determined that this image was from CSC eye, but Keras model judged from a Normal eye. b. Heatmep superimposed on OCT *en face* image. DL on the uniformity of the choroidal vessels and the area occupied by the choroidal vessels in the image. The active region is 74.24%.

### Deep learning and Heatmap

In both the NNC and Keras models, the *en face* images of 45 CSC eyes were used for training and 8 CSC eyes for validation. The *en face* images of 35 normal eyes were used for training and 12 for validation. The accuracy rate was 20 of 20 (100%) for NNC and 19 of 20 (95%) for Keras. The difference between the NNC and Keras models was not significant (*P* = 0.33).

The mean active region (± SD%) in the heatmap image generated by Grad-CAM was 12.5 ± 23.7% for CSC eyes which was significantly lower than the 79.8 ± 17.3% for normal eyes (*P* <0.01). One case of CSC (Figure 4) had a high value of 74.24%, and this case was incorrectly determined as normal in Keras. If that case was excluded, the mean active region in the CSCs eyes was very low at 3.7 ± 4.2%.

## Discussion

Our results showed that both of our DL models can accurately distinguish normal from CSC eyes by analyzing high-resolution choroidal vascular *en face* OCT images. The DL focused on the uniformity of the choroidal vessels and the area occupied by the choroidal vessels in the images.

At present, imaging diagnosis using DL in the medical field is rapidly developing and has become common in the fields of radiology and pathology.[34–37] In ophthalmology, it has also been shown to be useful in the imaging diagnosis and staging of various diseases such as diabetic retinopathy,[4–6] glaucoma,[7–9] age-related macular degeneration[10,11], and retinopathy of prematurity [12–14]. In the current study, we applied DL methods to evaluate the pattern of the choroidal vessels, the primary cause of CSC. The diagnosis of CSC is usually made by OCT and angiography in addition to fundus findings and cannot be determined by *en face* OCT alone. This study showed that DL can be used to diagnose typical CSC eyes with almost 100% accuracy based on the choroidal vascularity. A side benefit of DL is that the findings that were previously thought to be ambiguous may be defined with certainty by re-evaluating them from a DL perspective. In addition, it is expected that the use of DL will lead to new findings that have not been noticed by specialists.

We determined what characteristics were examined by DL of the Keras model using heatmaps with a Grad-CAM implementation. Until now, retinal specialists have observed the choroidal vascular characteristics in OCT *en face* images using choroidal vascular dilation, its flow pattern, and density. However, our results showed that DL may have focused on the uniformity of the choroidal vessels and were more evenly distributed in the images of normal eyes than in CSC eyes. It is true that the choroidal vessels are only visible in the upper or lower half of the image in some eyes with CSC. While we were focusing on the asymmetry of the choroidal vessel pattern, we were ignoring the fact that uniformity itself is a diagnostic marker. This unprecedented perspective may be an expected aspect of DL in the future.

There are limitations in this study including its retrospective nature, and it was performed on only a small number of cases. In addition, there was a significant difference in the ages in the normal and CSC groups. This is important because the choroid is thicker, and the density of the vessels is higher in younger individuals. These differences need to be considered, especially when the two groups are highly similar. However, this age difference is unlikely to have affected the DL because the results were almost completely classified using the DL. In addition, even though large choroidal vessels exist in the deeper layers, we have evaluated only one-half of the choroidal vascularity in the *en face* OCT images. However, we believe that this method has already been reported and is adequate to assess the medium and large vessels in the choroid with good accuracy. [25][38] The blood vessels information at the optic disc was excluded from the choroidal blood flow, while the artifacts due to large retinal vessels cannot be excluded. Because the luminal area in the retinal blood vessel is much smaller than in the choroidal blood vessel, it is generally believed that there is only a small influence of the retinal vessels. We also had cropped the *en face* images to exclude the influence of the optic disc in our analyses. Thus, these variables have been excluded in the analyses.

In conclusion, DL using 2 CNN models is able to accurately differentiate normal and CSC eyes from the *en face* OCT images of the choroidal vasculature alone. Previously, the classification mechanism was generally a black box, but according to a heatmap analysis using Keras with Grad-CAM implementation, DL does not evaluate the choroidal vessel dilation or flow pattern, it focuses on its uniformity and the area occupied by choroidal vessels in the image. It was interesting to see how these results differed from the focus of retinal specialists. In the future, physicians may not only be receiving assistance in diagnosis of DL, but they may also learn a new perspective from the process of reaching that diagnosis.

## Acknowledgments

The authors thank Professor Emeritus Duco Hamasaki of the Bascom Palmer Eye Institute, University of Miami, for his critical discussion and editing of the final manuscript.

